# A conserved family of immune effectors cleaves cellular ATP upon viral infection

**DOI:** 10.1101/2023.01.24.525353

**Authors:** Francois Rousset, Erez Yirmiya, Shahar Nesher, Alexander Brandis, Tevie Mehlman, Maxim Itkin, Sergey Malitsky, Adi Millman, Sarah Melamed, Rotem Sorek

## Abstract

During viral infection, cells can deploy immune strategies that deprive viruses of molecules essential for their replication. In this study, we report a family of immune effectors in bacteria which, in response to phage infection, degrade cellular ATP and dATP by cleaving the N-glycosidic bond between the adenine and sugar moieties. These ATP nucleosidase effectors are widely distributed within multiple bacterial defense systems including CBASS, prokaryotic argonautes and NLR-like proteins, and we show that degradation of (d)ATP during infection halts phage propagation and aborts infection. By analyzing homologs of the immune ATP nucleosidase domain, we discover and characterize Detocs, a new family of bacterial defense systems with a two-component phosphotransfer signaling architecture. The immune ATP nucleosidase domain is also encoded within a diverse set of eukaryotic proteins that have immune-like architectures, and we show biochemically that these eukaryotic homologs preserve the ATP nucleosidase activity. Our findings suggest that ATP and dATP degradation is a cell-autonomous innate immune strategy conserved across the tree of life.

## Introduction

The coevolution between viruses and their hosts has driven the emergence and diversification of numerous innate immune strategies in all domains of life. It was recently shown that multiple components of the cell-autonomous innate immune system of animals are also conserved in bacteria, where they function to protect against phage infection^1^. Such conserved immune components include the cGAS-STING pathway^2–4^, NOD-like receptors (NLRs)^5,6^, gasdermin-mediated pyroptosis^7^, Toll/interleukin-1 receptor (TIR) domain signaling^8,9^, the RNAi pathway^10,11^, SAMHD1^12^, viperins^13^, and others^14^. Based on these discoveries, it was shown that studying the immune system of bacteria can generate functional knowledge on previously unknown eukaryotic immune mechanisms^8,14,15^.

The human cGAS-STING pathway, which senses viral infection in human cells, was shown to be evolutionarily derived from a bacterial defense system called CBASS (cyclic oligonucleotide-based antiviral signaling system)^2–4^. Bacterial CBASS immunity relies on a cGAS/DncV-like nucleotidyltransferase (CD-NTase) enzyme that, upon sensing phage infection, produces a cyclic oligonucleotide molecule that binds and activates a cell-killing effector protein^16^. CBASS systems are found in 14% of sequenced bacterial and archaeal genomes and show substantial diversity in their oligonucleotide signals and effector functions^17,18^. CBASS effector proteins can exert their cell-killing function by degrading phage and host DNA^19–21^, by disrupting the cell membrane integrity^3,22^ or by depleting the essential molecule NAD^+ 4,23,24^.

A recent analysis of CBASS diversity in prokaryotic genomes revealed that ∼10% of type III CBASS systems encode an effector called Cap17, whose molecular function has remained unknown^18^ (**Fig. 1A**). In this study, we show that Cap17 is an ATP nucleosidase that cleaves ATP and dATP molecules into adenine and (deoxy)ribose-5’-triphosphate during phage infection, depriving the phage of these essential molecules and aborting the infection process. We further show that the Cap17 ATP nucleosidase domain is a distinct immunity domain widespread in multiple types of anti-phage systems, including prokaryotic argonautes, NLR-like proteins, and others. By studying operons of unknown function that contained the Cap17-like ATP nucleosidase domain, we discovered Detocs, a two-component signal transduction immune pathway that mediates ATP degradation upon sensing viral infection. Homologs of the Cap17 ATP nucleosidase domain are widespread in eukaryotes, from fungi to animals, where they are associated with various proteins with immune-like architecture. Our study defines a new immune function that is conserved across the tree of life.

**Figure 1.**
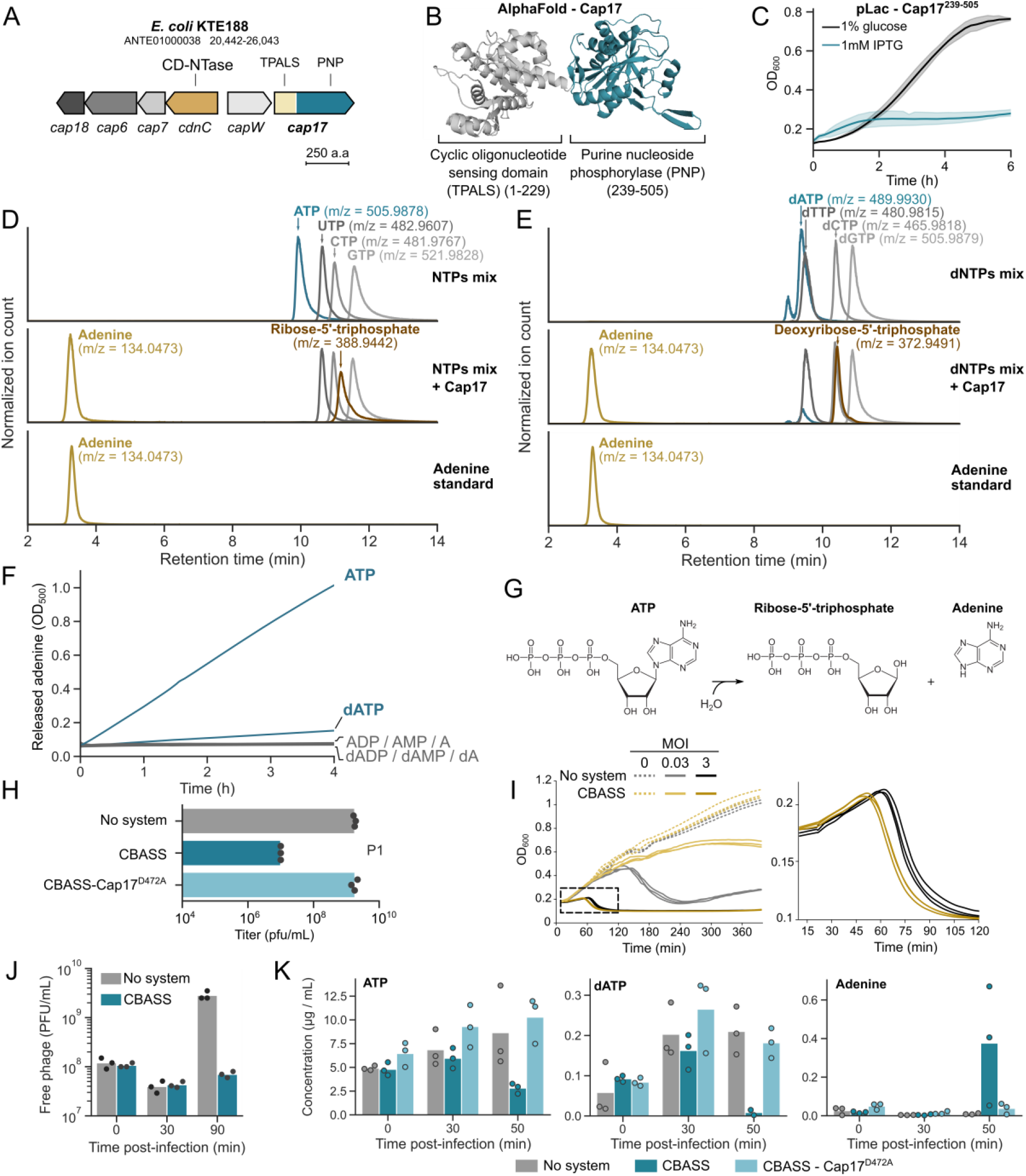
CBASS-associated Cap17 cleaves ATP molecules during phage infection. (A) A type III CBASS from *E. coli* KTE188 encodes the Cap17 effector (**Table S1**). Genbank accession and genome coordinates are displayed on top. Abbreviations: CD-NTase, cGAS/DncV-like nucleotidyltransferase; TPALS, TIR- and PNP-associating SLOG family; PNP, purine nucleoside phosphorylase. (B) Predicted AlphaFold structure of Cap17 from *E. coli* KTE188. (C) Growth curves of *E. coli* K-12 cells expressing the PNP domain from Cap17 (residues 239-505) from the pLac promoter under conditions repressing (1% glucose, black) or inducing expression (1 mM IPTG, blue). Curves show the mean of three replicates with the standard deviation shown as a shaded area. (D) LC-MS analysis of enzymatic reactions with NTPs in the absence (top) or presence (middle) of the Cap17 protein. Peak intensities of each compound were normalized across all samples in the *in vitro* assays. A synthetic adenine standard is shown (bottom). Ribose-5’-triphosphate was identified by MS/MS (**Fig. S2A**). (E) LC-MS analysis of enzymatic reactions with dNTPs in the absence (top) or presence (middle) of the Cap17 protein. Peak intensities of each compound were normalized across all samples in the *in vitro* assays. A synthetic adenine standard is shown (bottom). Deoxyribose-5’-triphosphate was identified by MS/MS (**Fig. S2B**). (F) Xanthine oxidase assays measuring adenine release from adenine-containing nucleotides by Cap17, as monitored by absorbance at 500 nm. (G) A schematic of the Cap17 enzymatic reaction. (H) Quantification of phage infection efficiency. Serial dilution plaque assay of phage P1 spotted on a lawn of *E. coli* K-12 cells expressing an empty vector (no system), a wild-type CBASS, or a mutated CBASS (CBASS-Cap17^D472A^). Bars represent the average of three replicates with individual data points overlaid. (I) Growth curves of *E. coli* K-12 cells expressing an empty vector or the CBASS system, infected by phage P1 at an MOI of 0.03 or 3 (or 0 for uninfected cells). The dashed area is magnified on the right. Three replicates are presented as individual curves. (J) Plaque-forming units of phage P1 sampled from the supernatant of *E. coli* K-12 cells expressing an empty vector or the CBASS system. T=0 represents initial phage titer prior to infection. Cells were infected at an MOI of 0.1. Bars represent the average of three replicates with individual data points overlaid. (K) Quantification of ATP, dATP and adenine in lysates derived from P1-infected *E. coli* cells expressing an empty vector (no system), a wild-type CBASS, or mutated CBASS (CBASS-Cap17^D472A^) by LC-MS. Bars represent the average of three replicates with individual data points overlaid.

## Results

### The CBASS effector Cap17 is an ATP nucleosidase

We set out to study the Cap17 protein from the type III CBASS of *E. coli* KTE188, a system that was previously shown to confer resistance against phages^25,26^ (**Fig. 1A**). Structural analysis of Cap17 via Alphafold2^27,28^ revealed a two-domain architecture with an N-terminal domain predicted to bind the cyclic oligonucleotide messenger (pfam PF18178) and a C-terminal domain of the PNP_UDP_1 family (pfam PF01048, hereafter called PNP) (**Fig. 1B, S1**). Expression of the PNP domain alone was toxic to *E. coli*, suggesting that it targets an essential host component (**Fig. 1C**). The PNP domain was originally described in diverse housekeeping enzymes involved in nucleotide salvage pathways, and is known to cleave the N-glycosidic bond in different nucleosides, separating the nucleobase from the sugar moiety^29^. We therefore suspected that the CBASS PNP effector might somehow manipulate cellular nucleotides or nucleosides as part of its anti-phage defensive activity.

We incubated purified Cap17 from *E. coli* KTE188 with a mixture of nucleotide triphosphates (NTPs) or deoxynucleotide triphosphate (dNTPs) and subjected the *in vitro* enzymatic reactions to liquid-chromatography followed by mass spectrometry (LC-MS). Incubation of purified Cap17 with NTPs revealed a specific consumption of ATP coupled with the formation of two products that we identified as adenine (m/z = 134.047) and ribose-5’-triphosphate (m/z = 388.944) (**Fig. 1D & S2A**). Levels of GTP, UTP and CTP were not affected by Cap17, suggesting specificity for ATP (**Fig. 1D**). Incubation of Cap17 with dNTPs also revealed consumption of dATP, coupled with the formation of adenine (m/z = 134.047) and deoxyribose-5’-triphosphate (m/z = 372.949) (**Fig. 1E & S2B**). These results suggest that Cap17 specifically cleaves the N-glycosidic bond of (d)ATP to release free adenine. Cap17 was unable to release adenine from DNA or RNA, suggesting that it is only active on free nucleotides (**Fig. S3**). To further assess the substrate specificity of Cap17, we measured its ability to release adenine from various substrates using a xanthine oxidase assay^30^ (**see Methods**). Cap17 activity was markedly higher on ATP than on dATP, and no activity was detected on other adenine-containing nucleotides or nucleosides (**Fig. 1F**). These results demonstrate that Cap17 is an ATP nucleosidase *in vitro* (**Fig. 1G**).

To investigate the immune function of Cap17 during phage infection *in vivo*, we cloned the *E. coli* KTE188 CBASS (**Fig. 1A, Table S1**) with its native promoter into *E. coli* K-12 MG1655 that naturally lacks CBASS, and subjected the resulting strain to phage infection. Phage P1 formed ∼100-fold fewer plaques on CBASS-expressing cells as compared to cells carrying a control vector, confirming the defensive capacity of this CBASS system^25,26^ (**Fig. 1H**). A single amino acid mutation in the predicted catalytic site of the Cap17 PNP domain abolished defense, suggesting that the enzymatic activity of Cap17 is essential for CBASS defense (**Fig. 1H**). When grown in liquid medium, CBASS-expressing cultures showed marked survival when infected with P1 phage at low multiplicity of infection (MOI), but the culture collapsed when infected with a high MOI, a hallmark of abortive infection in which infected cells die before the completion of the phage cycle^16,31^ (**Fig. 1I**). Concomitantly, infected cells did not produce phage progeny, confirming that this CBASS system prevents phage propagation (**Fig. 1J**).

Analysis of the metabolite content in lysates derived from infected cells showed a marked decrease in ATP levels in CBASS-expressing cells, as well as a complete elimination of dATP, 50 minutes from the onset of infection (**Fig. 1K**). At the same time, an accumulation of adenine was observed (**Fig. 1K**). These changes were not observed in a strain where Cap17 was mutated to disrupt the PNP active site (D472A) (**Fig. 1K**). Since dATP is substantially less abundant than ATP in *E. coli* cells, the weak activity of Cap17 on dATP compared to ATP (**Fig. 1F**) might still be sufficient to cause the observed depletion of dATP. Alternatively, dATP depletion could be a consequence of ATP degradation, given that cellular deoxynucleotides are synthesized from ribonucleotides^32^. A notable drop in the levels of ADP, dADP and AMP was also observed (**Fig. S4A**), but since those nucleotides are poor Cap17 substrates *in vitro* (**Fig 1F**), we suspect that their levels were affected as a secondary consequence of (d)ATP degradation.

In parallel with the depletion of ATP and dATP from infected CBASS-expressing cells, we observed an accumulation of the three other NTPs (GTP, CTP and UTP) and dNTPs (dGTP, dCTP and TTP) (**Fig. S4B-C**). This suggests that Cap17-mediated reduction in ATP and dATP levels disrupts the ability of the phage to produce nascent RNA and DNA chains respectively, resulting in the accumulation of unused non-adenosine (d)NTPs^12^. In support of this hypothesis, RNA and DNA sequencing revealed that the presence of CBASS reduced phage transcription and DNA replication (**Fig. S5A-B**). However, this reduction in phage transcription and DNA replication is likely insufficient to explain why CBASS-expressing cells disintegrate without producing viable phage progeny (**Fig. 1I-J**).

It was previously shown that exposure of phage-infected cells to chemicals that deplete cellular energy (“energy poisons” that disrupt the proton motive force) induces premature lysis^33^. This phenomenon, which was observed with multiple different phages, indicates that intracellular energy levels serve to regulate lysis timing by the phage lysis machinery^33^. We therefore hypothesize that ATP depletion by Cap17 causes premature lysis of infected cells (**Fig. 1I**). In agreement with this hypothesis, treatment of P1-infected control cells with the energy poison cyanide 3-chlorophenylhydrazone reduced lysis timing (**Fig. S5C**). Notably, overexpression of the PNP domain of Cap17 in uninfected cells caused a bacteriostatic effect but did not cause cell lysis (**Fig. 1C**), suggesting that the earlier culture collapse of P1-infected CBASS-expressing cells results from the combined effect of Cap17 activity together with phage factors (**Fig. 1I**).

Altogether, our results show that Cap17 is an immune effector whose activation leads to ATP and dATP degradation in response to phage infection, causing pleiotropic effects that prevent phage propagation.

### A specialized immune nucleosidase domain functions in multiple defense systems

Given that most proteins that encode PNP domains are housekeeping enzymes involved in nucleotide recycling pathways^29^, the observed antiviral role of the Cap17 PNP domain was puzzling. We therefore wondered whether the PNP domain of Cap17 represents a distinct subfamily that became specialized for an immune function. To examine this possibility, we analyzed a set of ∼38,000 bacterial and archaeal genomes and collected ∼110,000 proteins in which the PNP_UDP_1 protein domain (pfam PF01048) was detected. We then constructed a phylogenetic tree based on a sequence alignment of the PNP domains only (**see Methods**). PNP domains clustered into several distinct clades on the tree, with most clades containing PNPs from well-known housekeeping enzymes, including DeoD, XapA, Udp and MtnN (**Fig. 2A**). The PNP domain of Cap17, however, clustered in a distinct clade that did not include any of the known housekeeping proteins (**Fig. 2A**).

**Figure 2.**
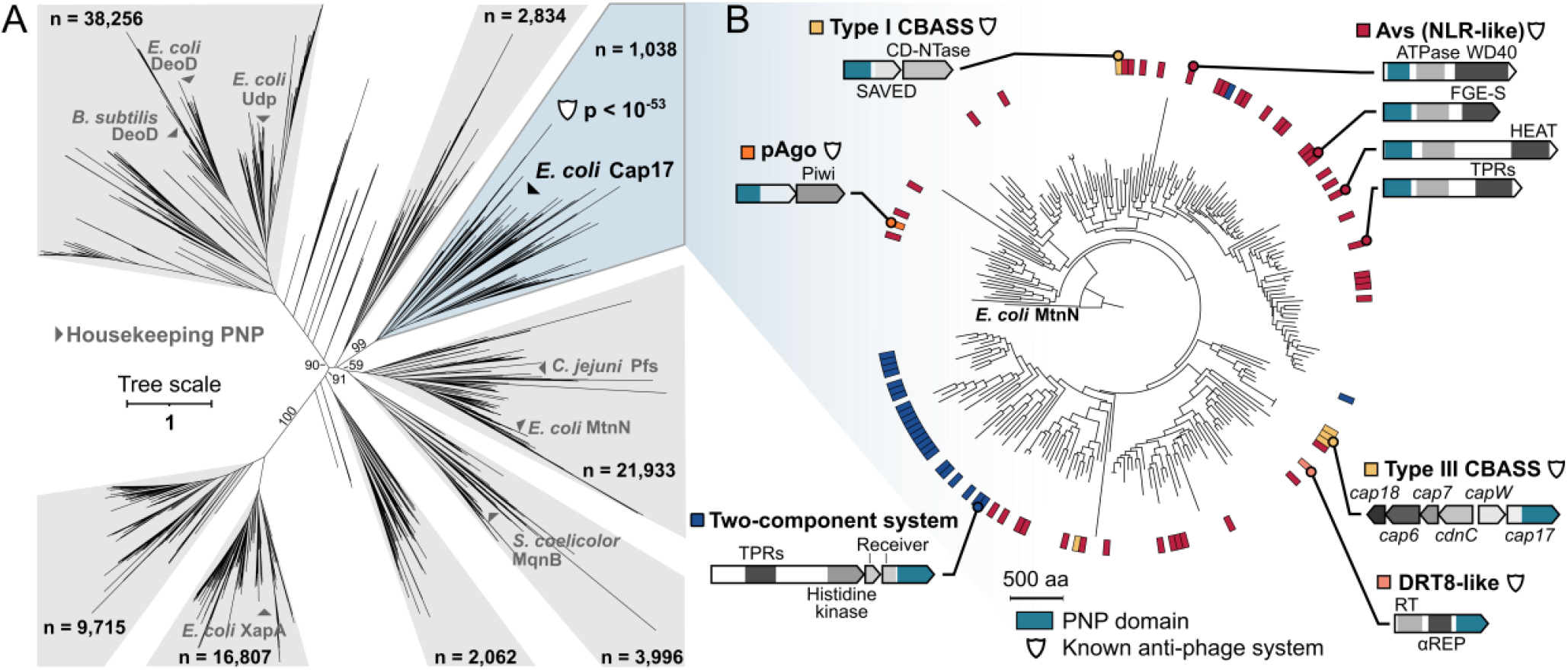
A clade of prokaryotic nucleosidases is associated with anti-phage systems. (A) Phylogenetic tree of prokaryotic PNP domains. Prokaryotic proteins were clustered based on sequence homology and clusters with homology to the PNP_UDP_1 domain (pfam PF01048) were identified. A representative sequence from each cluster was used to build the tree. The tree is based on the PNP domain only (**see Methods**). The total number of proteins represented by the representative sequences is shown for each clade. Ultrafast bootstrap values are shown for major branches^42^. Known housekeeping proteins are shown in grey. The p-value in the Cap17 clade represents the false discovery rate of a binomial test assessing the association of proteins from this clade with known anti-phage systems (**see Methods and Table S3**). (B) A detailed phylogenetic tree of the Cap17 PNP clade (marked blue in panel A). The PNP domain of *E. coli* MtnN was used as an outgroup. The colored ring represents known or predicted anti-phage systems harboring a PNP domain, with example systems shown as gene cassettes. Abbreviations: pAgo, prokaryotic argonaute; CD-NTase, cGAS/DncV-like nucleotidyltransferase; SAVED, SMODS-associated and fused to various effector domains; NLR, NOD-like receptor; FGE-S, Sulfatase-modifying factor enzyme 1; TPRs, Tetratricopeptide repeats; DRT8, Defensive Reverse-Transcriptase type 8.

We next examined the gene neighborhood of the 1,038 PNP-containing genes that were found in the Cap17 clade on the phylogenetic tree (**Fig. 2A**). This analysis showed that genes in this clade were very frequently found in the vicinity of known anti-phage systems in microbial genomes (corrected p-value < 10^−53^, **Fig. 2A, Table S3**), a metric that was previously shown to be a strong predictor of antiviral activity^34–37^. None of the other major clades on the PNP tree showed a statistically significant enrichment next to known defense systems (**Table S3**), suggesting that the Cap17 clade represents an evolutionarily derived PNP domain that was adapted for immune functions.

Inspection of the genes from the Cap17 clade showed that this domain is embedded in multiple known anti-phage defense systems (**Fig. 2B, Table S4**). In addition to its presence within CBASS systems, we found this domain in the N-terminus of diverse genes of the Avs family ^5,6^. These genes represent a large family of immune proteins that are the likely ancestors of human and plant NOD-like receptors (NLRs) and inflammasomes^5,6^. In prokaryotes, the C-terminal domain of these proteins senses infection signals, and the N-terminus contains an effector domain that leads to cell suicide once infection is sensed^5^. The presence of the PNP domain in the N-termini of multiple Avs proteins strongly suggests that it functions as the cell-targeting effector domain in these proteins. We also found the PNP domain in short prokaryotic argonaute (pAgo) systems, which sense phage infection via a pAgo protein and induce cell death via an associated effector protein^38–41^. Effector proteins of short pAgo systems are diverse, and include protein domains that disrupt membrane integrity or cleave cellular NAD^+ 38–41^. Again, the presence of the PNP domain as an effector of prokaryotic pAgo defense systems indicates that these pAgo systems inflict abortive infection by PNP-mediated ATP degradation (**Fig. 2B**). Finally, we recorded a PNP domain in a type 8 Defense-associated Reverse Transcriptase (DRT8), where it replaces the usual nuclease effector^43^. Altogether, our findings suggest that ATP nucleosidases are immune effectors in diverse antiviral systems in prokaryotes.

### A family of two-component systems defends by ATP degradation

Some of the PNP domains from the defensive clade appeared in operons not previously reported as defensive (**Table S4**). Specifically, we observed multiple cases in which this domain was embedded in an operon whose architecture bears strong similarity to bacterial two-component signal transduction systems^44^ (**Fig. 2B, Fig. 3A**). Such signal transduction systems are ubiquitous in prokaryotes and comprise a sensor kinase that typically senses an environmental signal through its N-terminal domain, triggering autophosphorylation of a conserved histidine residue on the C-terminal kinase domain^44^. The phosphate group is then transferred to a conserved aspartate on the N-terminal receiver domain of the second protein, called the “response regulator”. Phosphorylation of the receiver domain activates the C-terminal domain of the response regulator, usually a DNA-binding domain that regulates the expression of target genes^44^. The PNP-containing operon identified here included a sensor kinase protein and a response regulator protein in which the usual C-terminal DNA-binding domain was replaced by the PNP domain (**Fig. 3A**). Since PNP domains from the Cap17 clade were associated with multiple defense systems, we hypothesized that this two-component-like operon represents a previously unidentified anti-phage system.

**Figure 3.**
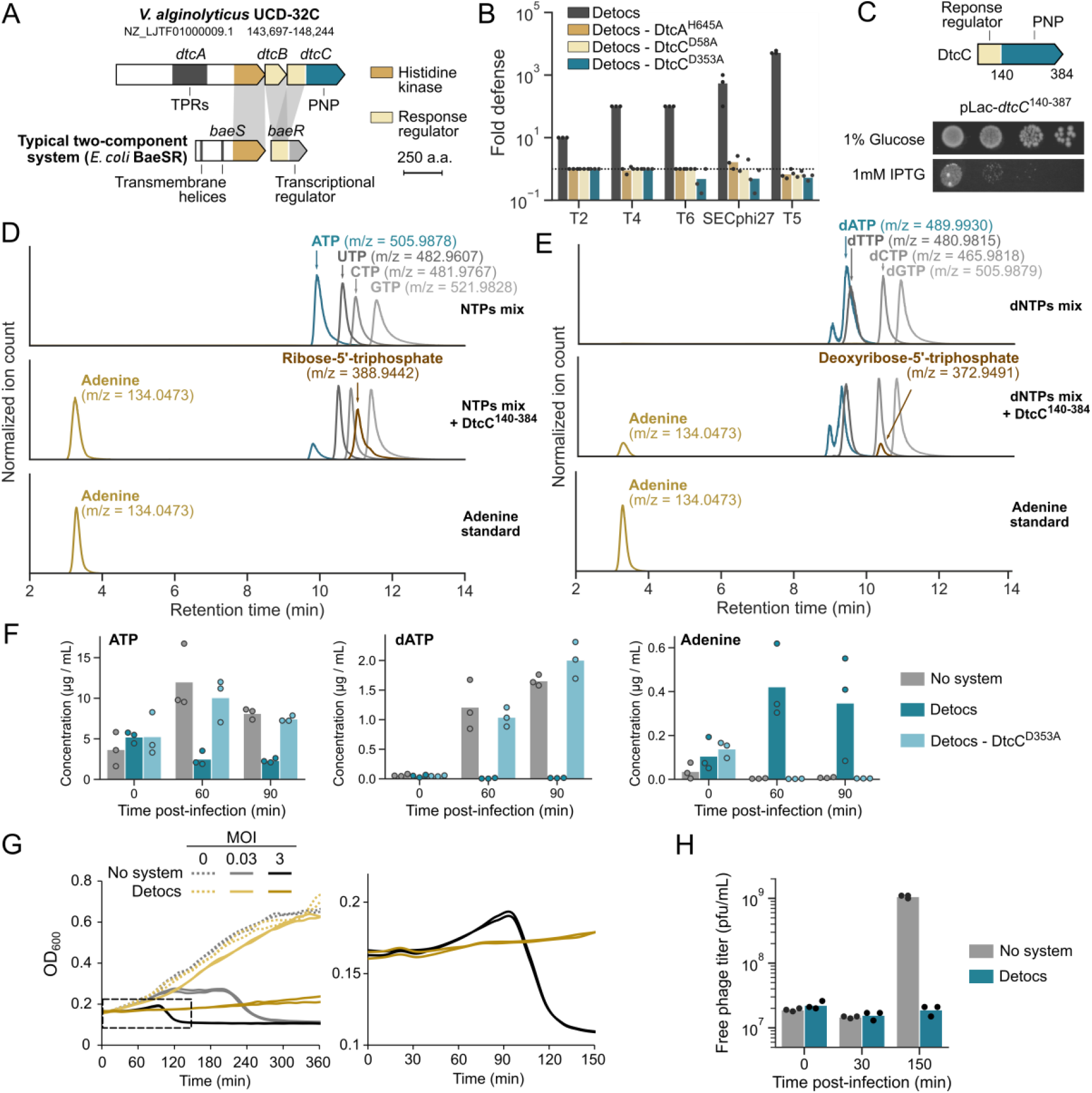
Detocs is a two-component system mediating ATP degradation upon viral infection. (A) A PNP-encoding genetic system from *V. alginolyticus* UCD-32C shares a similar architecture with typical two-component signal transduction systems. The *E. coli* BaeSR two-component system is depicted for comparison of domain architecture. TPRs, Tetratricopeptide repeats. (B) Detocs provides phage resistance. Serial dilution plaque assays were performed with phages infecting *E. coli* strains that expressed wild-type *V. alginolyticus* Detocs or Detocs mutated in the phosphate-binding histidine (Dtc^AH645A^), in the phosphate-receiving aspartate (DtcC^D58A^) or in the predicted catalytic site of the PNP domain (DtcC^D353A^), all expressed from the pBAD promoter. Fold defense was calculated as the ratio between phage plaque-forming units (PFUs) obtained on control cells and on defense-containing cells. Bars show the mean of three replicates with individual data points overlaid. (C) Ten-fold serial dilutions of *E. coli* cells expressing the PNP domain from DtcC (residues 140-387) from the pLac promoter were spotted on agar plates under conditions repressing (1% glucose) or inducing expression (1 mM ITPG). (D) LC-MS analysis of enzymatic reactions with NTPs in the absence (top) or presence (middle) of the PNP domain from DtcC (residues 140-387). A synthetic adenine standard is shown (bottom). (E) LC-MS analysis of enzymatic reactions with dNTPs in the absence (top) or presence (middle) of the PNP domain from DtcC (residues 140-387). A synthetic adenine standard is shown (bottom). (F) Quantification of ATP, dATP and adenine in lysates derived from T5-infected *E. coli* cells expressing an empty vector (no system), a wild-type Detocs, or mutated Detocs (Detocs-DtcC^D353A^). Bars represent the average of three replicates with individual data points overlaid. (G) Growth curves of *E. coli* cells expressing an empty vector or the Detocs system, infected by phage T5 at an MOI of 0.03 or 3 (or 0 for uninfected cells). The dashed area is magnified on the right. Three replicates are presented as individual curves. (H) Plaque-forming units of phage T5 sampled from the supernatant of *E. coli* cells expressing an empty vector or the Detocs system. T=0 represents initial phage titer prior to infection. Cells were infected at an MOI of 0.1. Bars represent the average of three replicates with individual data points overlaid.

To test this hypothesis, we cloned such an operon from *Vibrio alginolyticus* UCD-32C (**Fig. 3A, Table S1**) under the control of an inducible promoter into *E. coli* K-12 and infected the resulting strain with a diverse panel of phages. The system provided strong resistance against phages T2, T4, T5, T6 and SECphi27, a defense phenotype that required an intact PNP active site (**Fig. 3B**). Defense was abolished when the conserved histidine on the sensor kinase was replaced by alanine, as well as when the phosphate-receiving aspartate residue in the response regulator protein was mutated (**Fig. 3B**). These results suggest that autophosphorylation of the sensor kinase and phosphotransfer to the PNP-containing response regulator are necessary for the defensive activity of this system. We therefore named this system Detocs (<u>de</u>fensive tw<u>o</u>-<u>c</u>omponent <u>s</u>ystem). In addition to the sensor kinase (*dtcA*) and the response regulator (*dtcC*), the Detocs operon also contains a third gene (*dtcB*) whose function is studied below (**Fig. 3A**).

The PNP domain of DtcC from *V. alginolyticus* was highly toxic when expressed alone in *E. coli* (**Fig. 3C**). *In vitro* biochemical assays showed that the purified PNP domain can release adenine from ATP and to a lesser extent from dATP, but does not affect other NTPs or dNTPs (**Fig. 3D-E**). In agreement with this observed biochemical activity, analysis of lysates from T5-infected cells showed a depletion of ATP and dATP in Detocs-expressing cells correlated with an accumulation of adenine, but not when the PNP domain was mutated in the active site (**Fig. 3F, S6**). These results demonstrate that the PNP domain performs a similar function when found in different defense systems. Detocs triggered growth arrest in cells infected with high MOI of phage T5 (**Fig. 3G**) and infected cells did not release phage progeny (**Fig. 3H**), confirming an abortive infection mechanism. Notably, unlike in CBASS (**Fig. 1I**), growth arrest was not associated with cell lysis, suggesting that the energy depletion induced by Detocs does not cause early lysis during T5 infection^33^ (**Fig. 3G**).

While the genetic architecture of Detocs is similar to that of regulatory two-component systems, Detocs also encodes another protein, DtcB, with a standalone receiver domain that is not linked to any effector domain (**Fig. 3A**). We found that a point mutation in the receiving aspartate residue of DtcB rendered Detocs toxic to the cell and that this toxicity was dependent on a functional PNP domain (**Fig. S7**). We therefore propose that DtcB serves as a “buffer” protein that absorbs phosphate signals that result from inadvertent leaky activation of DtcA in the absence of phage infection, thus preventing autoimmunity. Future studies will be necessary to test this hypothesis.

Homology-based analyses showed that Detocs systems are found in hundreds of bacterial genomes from diverse phyla^45^ (**Fig. S8A-B**). While 80% of Detocs operons encode PNP effectors, in a minority of these operons the PNP is replaced by other domains known to function as cell-killing effectors in bacterial defense systems, including endonuclease and transmembrane-spanning domains (**Table S5**). We experimentally tested a Detocs operon with a transmembrane α/β hydrolase effector from *Enterobacter cloacae* JD6301, and found that it was able to efficiently protect *E. coli* against diverse phages (**Fig. S8C, Table S1**).

### ATP nucleosidases in eukaryotic innate immune proteins

Our findings demonstrate that the presence of the Cap17-like PNP domain within bacterial proteins of unknown function is predictive of immune activity of these proteins. This motivated us to ask whether Cap17-like ATP nucleosidases may function in eukaryotes as innate immune proteins, since multiple components of the innate immune system of eukaryotes were recently shown to be evolutionarily derived from bacterial defense systems^1^. To this end, we collected from Uniprot ∼10,000 eukaryotic proteins encoding the PNP_UDP_1 domain of pfam PF01048. We then reconstructed a phylogenetic tree that includes all detected PNP domains from both eukaryotes and prokaryotes (**Fig. 4**). PNP domains from housekeeping human proteins clustered with PNP domains of housekeeping bacterial proteins, but none of these were included in the defensive clade that contained the PNPs of CBASS and Detocs (**Fig. 4**). However, numerous eukaryotic proteins, whose functions were uncharacterized to date, clustered within the defensive PNP clade, hinting at the presence of an ATP nucleosidase immune activity in eukaryotes (**Fig. 4, Table S6**).

**Figure 4.**
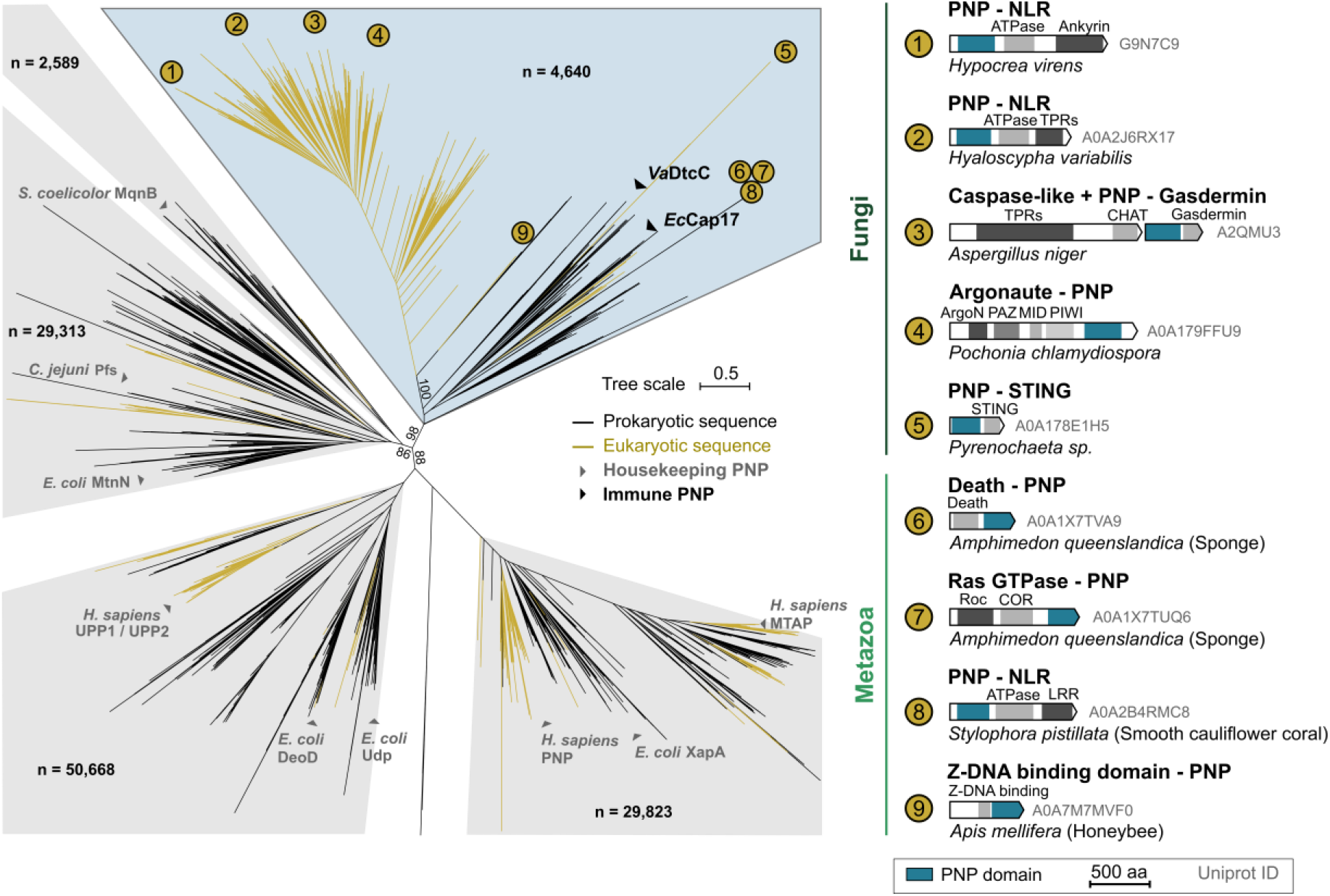
Diverse eukaryotic immune proteins carry Cap17-like PNP effector domains. Phylogenetic tree of prokaryotic and eukaryotic PNP domains. Prokaryotic and eukaryotic proteins were separately clustered based on sequence homology, and a representative sequence from each cluster was used to build the tree. Tree is based on the PNP domain only (**see Methods**). Ultrafast bootstrap values are shown for major branches42. Prokaryotic and eukaryotic PNP domains are shown as black and yellow branches, respectively. PNP domains from known housekeeping proteins are annotated in grey. The immune PNP domains of *V. alginolyticus* DtcC (VaDtcC) and of *E. coli* Cap17 (EcCap17) are in black. The total number of redundant protein sequences represented in each clade is depicted. The architecture of selected eukaryotic homologs is displayed on the right. Abbreviations: NLR, NOD-like receptor; TPRs, Tetratricopeptide repeats; CHAT, Caspase HetF Associated with TPRs; PAZ, Piwi Argonaute and Zwille; STING, Stimulator of Interferon Genes; Roc, Ras of Complex; COR, C-terminal of Roc; LRR: Leucine-rich repeats.

The domain architecture of eukaryotic proteins that cluster within the defensive PNP clade is strongly suggestive of immune functions (**Fig. 4, Table S6**). Specifically, there were hundreds of fungal and coral proteins with an NLR architecture, including a C-terminal sensing domain, a central nucleotide-binding domain and an N-terminal PNP domain^46,47^ (**Fig. 4**). NLR proteins are abundant pathogen sensors in animals and plants, and fungal NLRs were shown to participate in innate immunity by mediating programmed cell death upon allorecognition^48^, a phenomenon that restricts mycovirus transmission^49^. The presence of the PNP domain in eukaryotic NLRs suggests that these proteins cleave ATP molecules in response to pathogen sensing. Defensive PNP domains were also found in other proteins, combined with additional domains that very strongly indicate an immune function. These included a PNP domain fused to a STING domain in the fungus *Pyrenochaeta*, suggesting that the PNP domain is activated in response to cGAS-like signaling^4,45^; a PNP domain fused to an argonaute protein in *Pochonia chlamydiospora*, suggesting that PNP activity might supplement an RNA interference response^50^; a Gasdermin protein fused to a PNP domain and linked to a nearby caspase-like protease in *Penicillium glabrum*, suggesting that this protein might be activated by proteolytic cleavage into a double-edged effector^7,51,52^; and fusions between PNP and Death domains which participate in the assembly of inflammasomes^53,54^, for example in the sponge *Amphimedon queenslandica* (**Fig. 4**).

Proteins with the Cap17-like PNP domain were also found in arthropods and nematodes, including in the well-studied model organisms *Drosophila melanogaster* (Uniprot accession: Q8MRM6*), Apis mellifera* (Uniprot accession: A0A7M7MVF0) and *Caenorhabditis elegans* (Uniprot accession: H2L0A5) (**Fig. 4, Table S6**). In addition to the PNP domain, these proteins also include a domain homologous to the Zα domain found in immune proteins such as human ZBP1 and ADAR1 where this domain senses foreign nucleic acids with Z-DNA or Z-RNA configuration^55,56^. It is tempting to speculate that these PNP-containing proteins in arthropods and nematodes degrade (d)ATP upon recognition of foreign nucleic acids.

To test whether the PNP domains of eukaryotic immune-like proteins might mediate (d)ATP degradation, we cloned a PNP domain from an NLR-like protein from the fungus *Hyaloscypha variabilis* and a Death-domain-associated PNP domain from the sponge *A. queenslandica* (**Fig. 5A**,**B**). Both domains were toxic when expressed in *E. coli* (**Fig. 5C**,**D**) and purified domains showed preference for ATP over dATP while being poorly active on other (deoxy)adenylate nucleotides (**Fig. 5E**,**F**). By incubating the PNP domain of *A. queenslandica* with pools of NTPs and dNTPs, we verified that the eukaryotic domain was active only against ATP and dATP (**Fig. 5G, H**). These data suggest that the function of this domain is preserved from prokaryotes to eukaryotes. Taken together, the *in vitro* activity of eukaryotic Cap17-like PNP domains, in addition to their presence within proteins with a typical immune architecture, strongly suggest that ATP degradation is a cell-autonomous innate immune strategy in both prokaryotes and eukaryotes.

**Figure 5.**
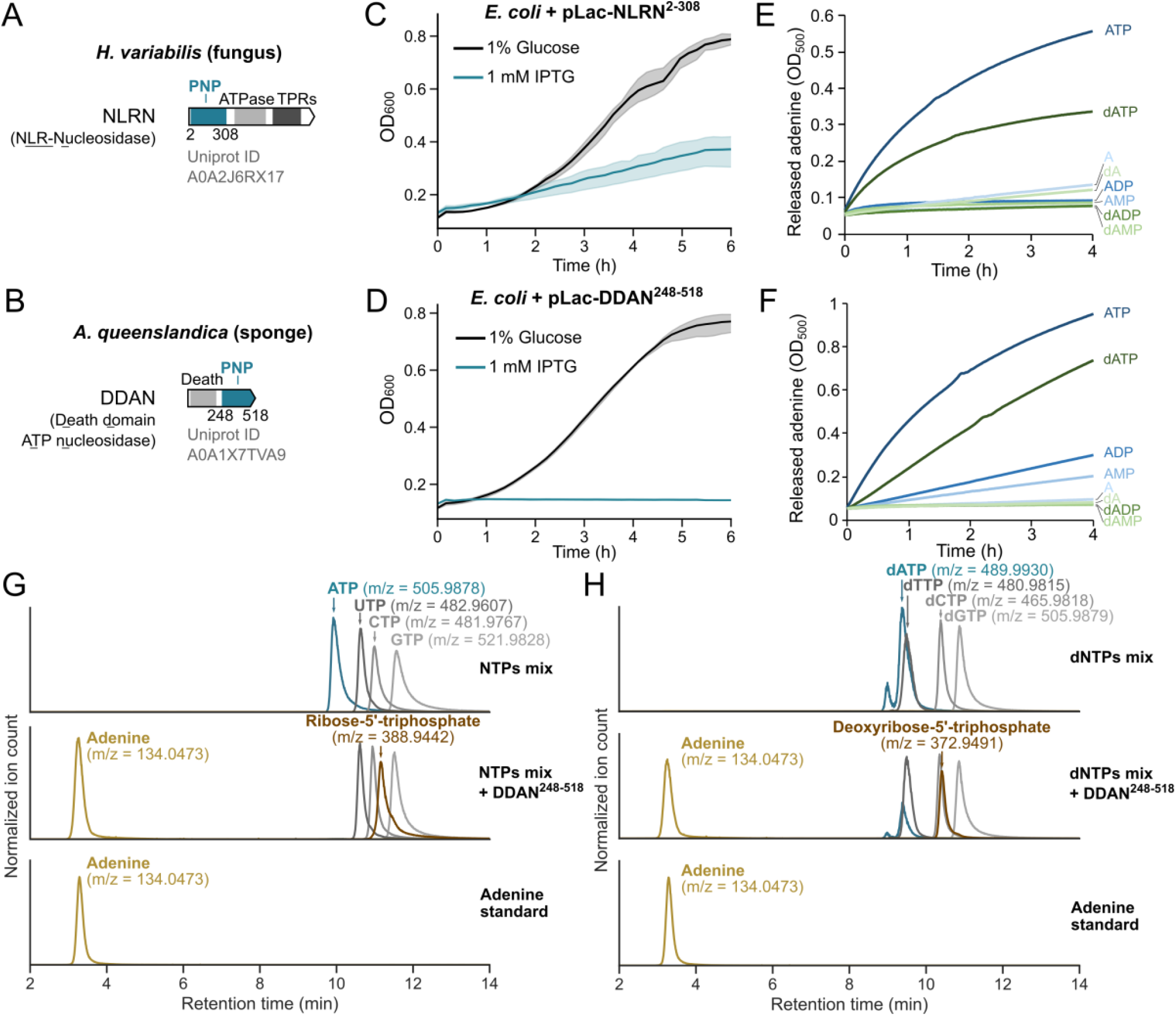
PNP domains associated with eukaryotic immune proteins have ATP nucleosidase activity. (A) The NLR-like protein NLRN from the fungus *Hyaloscypha variabilis* harbors an N-terminal PNP domain (residues 2 to 308). TPRs, Tetratricopeptide repeats. (B) The death domain-containing protein DDAN from the sponge *Amphimedon queenslandica* harbors a C-terminal PNP domain (residues 248 to 518). (C-D) Growth curves of *E. coli* cells expressing the PNP domain from (C) NLRN or (D) DDAN from the pLac promoter in conditions repressing expression (1% glucose, black) or inducing expression (1 mM ITPG, blue). Curves show the mean of three replicates with the standard deviation shown as a shaded area. (E-F) Xanthine oxidase assays measuring adenine release from adenine-containing nucleotides by (E) the PNP domain of NLRN or (F) the PNP domain of DDAN, as monitored by absorbance at 500 nm. (G) LC-MS analysis of enzymatic reactions with NTPs in the absence (top) or presence (middle) of the PNP domain from DDAN. A synthetic adenine standard is shown (bottom). (H) LC-MS analysis of enzymatic reactions with dNTPs in the absence (top) or presence (middle) of the PNP domain from DDAN. A synthetic adenine standard is shown (bottom).

## Discussion

In this study, we describe a new immune strategy mediated by a family of enzymatic effectors that degrade (d)ATP in response to viral infection. This discovery expands the repertoire of immune domains known to target essential metabolites to limit viral replication, including the TIR^4,23,24,38,57^, SIR2^8,40,41^ and SEFIR^37^ domains that degrade NAD^+^, and dCTP deaminases^12^, dGTPases^12^ and SAMHD1^58^ that degrade deoxynucleotides. The central role of ATP makes it an ideal target for degradation in order to simultaneously inhibit multiple cellular processes that are essential for viral propagation. The Cap17 ATP nucleosidase activity separates the adenine base from the sugar moiety, generating free adenine and (deoxy)ribose-5’-triphosphate, a molecule not naturally synthesized by *E. coli* cells to our knowledge. Therefore, it is unlikely that the molecules generated by Cap17 could be efficiently used by cellular pathways to rebuild ATP.

Interestingly, ATP and dATP degradation by Cap17 leads to cell lysis during phage P1 infection (**Fig. 1I**), while the expression of ATP nucleosidase domains alone leads to growth arrest but does not induce cell lysis (**Fig. 1C**). The phenomenon of early cell lysis following energy depletion was also observed in the context of SIR2, TIR and SEFIR immunity, in which NAD^+^ degradation caused the cells to lyse prematurely after infection by some phages^8,37,41,57^. Our data may suggest that energy depletion during infection not only deprives phages of the energy necessary for their propagation, but also serves to dysregulate their lysis machinery^33^, forcing it to prematurely lyse the host cell prior to the completion of the infection cycle.

By analyzing homologs of the Cap17 ATP nucleosidase domain, we discovered Detocs, a family of defensive two-component signal transduction pathways. Two-component signal transduction systems are extremely abundant in bacteria, where they frequently mediate transcriptional regulation and chemotaxis in response to specific environmental signals^44^. While some two-component systems were shown to regulate the expression of anti-phage systems^59,60^, our data show for the first time that two-component signal transduction pathways were evolutionarily adopted as *bona fide* anti-phage defense systems. The genetic architecture of Detocs supports a mechanism in which phage infection is sensed by the DtcA kinase, which autophosphorylates and transmits the phosphate signal to the DtcC effector to activate it, as previously hypothesized based on a bioinformatic analysis^45^. The system also contains DtcB, an additional protein encoding a single receiver domain, which we hypothesize prevents autoimmunity by scavenging phosphate signals resulting from unintended activation of DtcA. Remarkably, the well-known modularity of two-component systems^44^ is preserved in Detocs, in which the ATP nucleosidase effector domain can be replaced by other effector types (**Fig. S8**), illustrating how swapping effector domains can expand the functional diversity of anti-phage systems.

Our evolutionary analyses suggest that immune ATP nucleosidases form an ancient subfamily within the PNP domain family. Based on the topology of the phylogenetic tree, these immune ATP nucleosidases likely emerged in bacteria and were acquired by eukaryotes, possibly on multiple occasions (**Fig. 4**). Our data show that eukaryotic domains of this subfamily preserve the ATP and dATP nucleosidase activities *in vitro*, and are frequently found in proteins that are very likely to be involved in innate immunity based on their domain architectures. Interestingly, ATP N-glycosidase enzymatic activity has been observed in a sponge lysate^61,62^, but the functional role of this activity remains unknown. Further studies will be necessary to investigate the immune functions of Cap17-like ATP nucleosidases in eukaryotes. Beyond the Cap17-like PNP subfamily, housekeeping PNP enzymes from nucleotide salvage pathways have been shown to indirectly regulate innate immune functions in humans and in *C. elegans* by affecting nucleoside concentrations^63,64^, illustrating the broad importance of the PNP protein family in the manipulation of cellular nucleotides in immune processes.

In recent years, multiple eukaryotic innate immunity pathways have been shown to have a deep evolutionary origin in prokaryotes^1,14^. Many of the discoveries showing that bacterial defense systems function similarly to their eukaryotic counterparts were enabled by translating prior knowledge of eukaryotic immunity into the prokaryotic world. However, the recent expansion of the known repertoire of anti-phage systems now provides the opportunity to apply the reverse reasoning, *i*.*e*. to translate our knowledge of prokaryotic immunity into the eukaryotic world^8,14,15,37^. Our discovery of a new immune effector domain that is conserved across the tree of life presents an opportunity to unravel previously unknown functions within the innate immune system of eukaryotes based on bacterial defense against phages.

## Acknowledgement

We thank Yao Li, Hunter Toyoda and Philip Kranzusch for scientific insights, Emily Troemel and Vladimir Lažetić for useful discussion on eukaryotic PNP domains, and all members of the Sorek lab for critical comments on the manuscript. F.R was supported by the Clore Foundation postdoctoral fellowship and by the Dean of Faculty fellowship from the Weizmann Institute of Science. R.S. was supported, in part, by the European Research Council (grant ERC-AdG GA 101018520), Israel Science Foundation (MAPATS grant 2720/22), the Deutsche Forschungsgemeinschaft (SPP 2330, grant 464312965), the Ernest and Bonnie Beutler Research Program of Excellence in Genomic Medicine, and the Knell Family Center for Microbiology. E.Y. is supported by the Israeli Council for Higher Education (CHE) via the Weizmann Data Science Research Center. A.M. was supported by a fellowship from the Ariane de Rothschild Women Doctoral Program and, in part, by the Israeli Council for Higher Education via the Weizmann Data Science Research Center. M.I. and S.Ma. are supported by the Vera and John Schwartz Family Center for Metabolic Biology.

## Author contributions

F.R., E.Y. and R.S. conceptualized the project. F.R., S.N. and S.Me. performed genetic analyses and plaque assays. F.R. performed protein purifications and biochemical assays. A.B., T.M., M.I. and S.Ma. performed LC-MS experiments. F.R. and E.Y. performed evolutionary analyses. F.R. and A.M. detected Detocs in bacterial genomes. F.R. and R.S wrote the manuscript.

## Methods

### Strains and media

*E. coli* NEB 5-alpha (New England Biolabs) was used as a cloning strain. *E. coli* K-12 MG1655 was used for phage-related experiments while *E. coli* BL21(DE3) was used for protein purifications. *Bacillus subtilis* BEST7003 was kindly provided M. Itaya. Unless mentioned otherwise, cells were grown in MMB (LB + 0.1 mM MnCl^2^ + 5 mM MgCl^2^, with or without 1.5% agar) with the appropriate antibiotics: ampicillin (100 µg/mL), chloramphenicol (30 µg/mL) or kanamycin (50 µg/mL).

### Plasmid construction

The type III CBASS system from *E. coli* KTE188 (Genbank: ANTE01000038)^26^ was synthesized by Genscript together with its native promoter and terminator sequences and cloned into plasmid pSG1 ^35^, yielding pSG1-CBASS (**Table S1**). Detocs systems from *V. alginolyticus* UCD-32C (Genbank: NZ_LJTF01000009.1) and from *Enterobacter cloacae* JD6301 (Genbank: JDWH01000005.1) were synthesized by Twist Bioscience as gene fragments and assembled into the pBAD plasmid^13^ under the arabinose-inducible pBAD promoter by Gibson assembly (New England Biolabs) (**Table S1**). Point mutations were introduced using the KLD Enzyme mix (New England Biolabs) (**Table S7**) and were verified by whole-plasmid sequencing (Plasmidsaurus). PNP domains from *Hyaloscypha variabilis* and *Amphimedon queenslandica* were synthesized by Twist Bioscience as gene fragments. All PNP domains tested for toxicity were cloned into the pBbA6c vector^65^ (Addgene plasmid # 35290) under the IPTG-inducible pLac promoter.

### Toxicity assays

To measure the toxicity of PNP domains, overnight cultures of *E. coli* K-12 MG1655 cells expressing the PNP domain from *E. coli* Cap17, *V. alginolyticus* DtcC, *H. variabilis* NLRN or *A. queenslandica* DDAN on the pBbA6c vector were diluted 100-fold in MMB + chloramphenicol supplemented with 1% glucose or 1 mM IPTG. Growth was followed by OD_600_ measurement every 10 min on a Tecan Infinite200 plate reader.

### Protein purification

DNA sequences encoding the proteins to purify were cloned onto the pET28-SUMO vector^15^ by Gibson assembly (**Table S8**). Plasmids were transformed into *E. coli* BL21(DE3) and clones were selected on LB medium supplemented with kanamycin. Overnight cultures of each resulting strain were grown in LB supplemented with kanamycin to OD_600_ of ∼0.8-1 and protein expression was induced by the addition of 0.5 mM IPTG (Inalco). Cultures were further incubated for 20 h at 16°C in the case of Cap17 or for 3 h at 37°C in the case of the PNP domains from *V. alginolyticus, H. variabilis* and *A. queenslandica*. Cells were harvested by centrifugation (5,000 g – 12 min) and resuspended in wash buffer (20 mM sodium phosphate buffer pH 7.4, 300 mM NaCl, 20 mM imidazole, 0.05% Tween20 supplemented with cOmplete Protease Inhibitor Cocktail tablet (Roche)). Resuspended cells were then transferred to a FastPrep Lysing Matrix B 2 ml tube (MP Biomedicals) and lysed using a FastPrep bead beater for 40 s at 6 m.s^−1^. After centrifugation (12,000 g – 10 min), the supernatant was bound to NEBExpress Ni-NTA Magnetic Beads (NEB) for 1 h at 4°C with rotation. Beads were washed 3 times with wash buffer and once with Ulp1 buffer (40 mM Tris-HCl pH 7.5, 250 mM NaCl, 250 mM sucrose, 2 mM MgCl^2^). The His-SUMO tag was cleaved on beads using Ulp1 protease produced in-house in Ulp1 buffer for 2 h at 25°C. Cleaved products were collected from the suspension and analyzed by SDS-PAGE and Coomassie staining. Proteins were flash-frozen and stored at -80°C until use.

### *In vitro* assays and LC-MS polar metabolite analysis

The substrate specificity of purified enzymes was assessed by incubating each of them with either a mix of NTPs (Ambion AM1322) or a mix of dNTPs (New England Biolabs N0447L) (final concentration: 100 μM each). Cap17 was also incubated with 50 ng/μL of lambda DNA (NEB #N3011S) or control RNA (Ambion MicrobExpress) to measure a possible release of adenine from nucleic acids. All reactions were performed in triplicates in a volume of 100 μL, in 100 mM sodium phosphate buffer (pH 7.4) with 500 nM enzymes for 1 h at 37°C, except for reactions with the PNP domain from *H. variabilis* which were performed with 1 μM enzyme for 4 h at 37°C. Before the injection into the LC-MS, samples were centrifuged twice (13,000 rpm) to remove possible precipitants, and transferred to an HPLC vial. Samples were analyzed as described previously^66^ with minor modifications described below. Briefly, analysis was performed using Acquity I class UPLC System combined with mass spectrometer Q Exactive Plus Orbitrap™ (Thermo Fisher Scientific), which was operated in a negative ionization mode. The MS spectra were acquired with 70.000 resolution, scan range of 100 – 800 m/z. For the identification of the compounds, we used a data-dependent acquisition, top 5 method. The LC separation was done using the SeQuant Zic-pHilic (150 mm × 2.1 mm) with the SeQuant guard column (20 mm × 2.1 mm) (Merck). The mobile phase B was acetonitrile and the mobile phase A was 20 mM ammonium carbonate with 0.1% ammonia hydroxide in a 80:20 solution (v/v) of double-distilled water and acetonitrile. The flow rate was kept at 200 μl.min^−1^, and the gradient was as follows: 75% of B (0-2min), decreased to 25% of B (2-14 min), 25% of B (14-18min), increased to 75% of B (18-19 min), 75% of B (19-23 min). Adenine was identified using a synthetic standard run (Sigma A2786). Data were analyzed using Mzmine 2.53 ^67^. For each compound, peak intensities were normalized across all samples.

### Xanthine oxidase assays

The substrate specificity of PNP enzymes on adenylate nucleotides was refined using a xanthine oxidase assay^30^. Briefly, the oxidation of released adenine to 2,8-dihydroadenine by xanthine oxidase is coupled to the reduction of two iodonitrotetrazolium chloride (INT) molecules into formazan which absorbs at 500 nm. Reactions were performed in 100 μL with 1 mM INT (Sigma I8377), 0.035 units of xanthine oxidase (Roche 10110434001) and 200 μM of each substrate (ATP, ADP, AMP, adenosine, dATP, dADP, dAMP and deoxyadenosine) in 100 mM sodium phosphate buffer (pH 7.4). Reactions were started by the addition of purified PNP enzymes to a final concentration of 200 nM, except for the PNP domain from *H. variabilis* which was added to a final concentration of 1 μM. Absorbance at 500 nm was followed on a Tecan Infinite200 plate reader.

### Plaque assays

All phages used in this study are listed in **Table S2**. Phages were amplified from single plaques in liquid cultures of *E. coli* K-12 MG1655 at 37°C in MMB medium until culture collapse. Cultures were centrifuged (4,000 g x 10 min) and the supernatants were passed through 0.2 µm filters.

Plaque assays were performed as previously described^68^. *E. coli* K-12 MG1655 carrying anti-phage systems (pSG1-CBASS or pBAD-Detocs) or control plasmids (pSG1 or pBAD) were grown overnight at 37°C in MMB medium supplemented with ampicillin. Then, 300 μL of each culture was mixed with 30 mL of molten MMB + 0.5% agar supplemented with 0.2% arabinose in the case of pBAD and pBAD-Detocs strains. The mixture was poured on 12 × 12 cm plates and left to dry for 1 h. Tenfold dilutions of phages were prepared in MMB and 10 µL of each dilution was dropped onto the plates. Plates were incubated overnight at 37°C (pSG1 and pSG1-CBASS) or 25°C (pBAD and pBAD-Detocs). Plaque-forming units (PFUs) were counted the next day.

### Liquid infection assays

Overnight cultures of *E. coli* K-12 MG1655 carrying anti-phage systems (pSG1-CBASS or pBAD-Detocs) or control plasmids (pSG1 or pBAD) were diluted 100-fold in MMB-ampicillin with (pBAD and pBAD-Detocs) or without (pSG1 and pSG1-CBASS) 0.2% arabinose and grown to an optical density at 600nm (OD_600_) of 0.3. Then, 190 µL of cultures were transferred to a 96-well plate and infected with phage P1 at 37°C (pSG1 and pSG1-CBASS) or with T5 at 25°C (pBAD and pBAD-Detocs) at various MOIs. Growth was followed by OD_600_ measurement every 5 min on a Tecan Infinite200 plate reader.

Propagation of phage P1 and T5 on control and defense-expressing cells was measured by infecting exponential cultures at OD_600_ ∼0.3 with P1 or T5 at MOI of ∼0.1. The infected cultures were sampled at the indicated time points, spun down (14,000 g – 1 min) and the free phage titer was assessed in the supernatant by plaque assays on *E. coli* K-12 MG1655 as described above.

### Cell lysate preparation

For CBASS, overnight cultures of *E. coli* K-12 MG1655 carrying pSG1, pSG1-CBASS or pSG1-CBASS-Cap17^D472A^ were diluted 100-fold in 200 mL of MMB + ampicillin and grown to OD_600_ ∼0.3 at 37°C with shaking (200 rpm). Then, 50 mL of cultures were sampled (Time 0) and the remaining cultures were infected with ∼10^11^ pfus of phage P1 (MOI ∼3). After 30 min and 50 min, 50 mL of the culture were sampled. For Detocs, sampling was performed similarly with phage T5 and the following exceptions: (i) MMB + ampicillin was supplemented with 0.2% arabinose, (ii) cultures were incubated at 25°C with shaking (200 rpm), (iii) cultures were sampled 60 min and 90 min post-infection. Following sampling, each sample was centrifuged (4,000 g – 7 min) at 4°C and cell pellets were flash-frozen in a bath of ethanol in dry ice. All pellets were thawed on ice, resuspended in 600 μL of cold 100 mM sodium phosphate buffer (pH 7.4), transferred to FastPrep Lysing Matrix B 2-ml tubes (MP Biomedicals, catalog no. 116911100) and lysed using a FastPrep bead beater for 40 s at 6 m.s^-1^. Samples were centrifuged (12,000 g – 10 min) and supernatants were passed through 3 kDa Amicon Ultra-0.5 Centrifugal Filter Units (Merck Millipore, catalogue no. UFC500396) for 45 min at 12,000 g at 4°C. Filtrates were used for LC-MS analysis for nucleotide quantification.

### Quantification of nucleotides by LC-MS

Quantification of metabolites was carried out using an Acquity I-class UPLC system coupled to Xevo TQ-S triple quadrupole mass spectrometer (Waters, US). The UPLC was performed using an Atlantis Premier BEH Z-HILIC column (2.1 × 100 mm, 1.7 um; Waters). Mobile phase A was 80% of 20mM ammonium carbonate, pH 9.25 in acetonitrile and mobile phase B was acetonitrile. The flow rate was kept at 300 μl.min^-1^ consisting of a 0.8 min hold at 80% B, followed by a linear gradient decrease to 25% B during 4.6 min. The column temperature was set at 35°C, and the injection volume was 3 µl. An electrospray ionization interface was used as an ionization source. Analysis was performed in positive ionization mode. Metabolites (d)GTP, (d)ATP, TTP/UTP, (d)CTP, (d)ADP, (d)AMP, (d)A, and adenine were detected using multiple-reaction monitoring, using argon as the collision gas. Quantification was made using a standard curve in 0.001–5 µg.mL^-1^ concentration range. ^13^C_10_-ATP (Sigma 710695) and ^15^N_5_-AMP (Sigma 662658) were added to standards and samples as internal standards to get 1 and 0.5 µM, respectively. TargetLynx (Waters) was used for data analysis.

### DNA and RNA sequencing

Overnight cultures of *E. coli* K-12 MG1655 carrying pSG1 or pSG1-CBASS were diluted 100-fold in 4 mL of MMB + ampicillin and grown at 37°C. When OD_600_ reached ∼0.3, cells were infected with phage P1 at an MOI of ∼3 and further incubated at 37°C. Before infection (time 0), and 30 min and 50 min after infection, 1 mL of culture was collected and added to 10 µL of pre-chilled overnight culture of *Bacillus subtilis* BEST7003 used as a spiked-in control. Samples were centrifuged at 4°C (16,000 g – 30 s) and pellets were flash-frozen in a bath of ethanol in dry ice. The remaining volume was infected with phage P1 at an MOI of ∼3 and further incubated at 37°C. Sample collection was repeated 30 min and 50 min after infection. For DNA sequencing, total DNA was extracted from pellets using the QIAGEN DNeasy blood and tissue kit (cat. #69504). Libraries were prepared for Illumina sequencing using a modified Nextera protocol as previously described^69^. For RNA sequencing, pellets were resuspended in 100 μl of 2 mg.ml^−1^ lysozyme in 10 mM Tris-HCl and 1 mM EDTA pH 8.0 and incubated at room temperature for 3 min. One milliliter of TRI-reagent (MRC) was added and samples were vortexed for 10 s before the addition of 200 μl chloroform. Following another vortexing step, the samples were left at room temperature for 5 min and centrifuged (16,000 g – 10min) at 4 °C. The upper phase was added to 500 μl of ice-cold isopropanol and samples were incubated at -20 °C for 1 h for RNA precipitation. Samples were then centrifuged (16,000-30min) at 4 °C and the RNA pellet was washed twice with ice-cold 70% ethanol, air-dried and resuspended in 30 μl water. RNA levels were measured using Nanodrop. All RNA samples were treated with TURBO DNase following the manufacturer’s instructions (Life technologies, AM2238). Ribosomal RNA depletion and RNA-seq libraries were prepared as previously described^70^, except that all reaction volumes were reduced by a factor of 4. All libraries were sequenced on an Illumina NextSeq500 sequencer. Reads were aligned to the genomes of *E. coli* K-12 MG1655 (Genbank NC_000913), *B. subtilis* BEST7003 (Genbank AP012496.1) and phage P1 (Genbank NC_005856.1) as described previously^70^. Read counts for each sample were normalized to the number of reads mapping the *B. subtilis* BEST7003 genome.

### Phylogenetic analysis of the PNP domain family

Protein sequences of all genes in 38,167 bacterial and archaeal genomes were downloaded from the Integrated Microbial Genomes (IMG) database^71^ in October 2017. Proteins with a hit to pfam PF01048 were collected and bacterial and archaeal sequences were separated. Proteins from each group were filtered for redundancy using the ‘clusthash’ option of MMseqs2 (release 12-113e3)^72^ using the ‘–min-seq-id 0.9’ parameter and then clustered using the ‘cluster’ option with the ‘--single-step-clustering’ flag. For each cluster, the representative sequence defined by MMseqs2 was collected and searched against the Pfam database^73^ using hhsearch^74^ with default parameters. Cluster representatives with a hit to pfam PF01048 with an e-value equal to or lower than 0.01 and spanning at least 100 residues were selected and the sequence of the PNP domain was extracted. All PNP domains from selected cluster representatives were aligned using Clustal-omega version 1.2.4^75^ with default parameters. The resulting multiple-sequence alignment was used to compute a tree using IQtree version 1.6.5^76^ using parameters -m LG -nt 1 -nm 5000. Node support was computed using 1000 iterations of the ultrafast bootstrap function in IQtree (option -bb 1000)^42^. All trees were visualized with iTOL^77^.

To find clades of PNP-containing genes that are significantly associated with defense systems (**Fig. 2A**), we collected all non-redundant PNP-containing genes and ran DefenseFinder v1.2^17^ on their genomes of origin. Then, for each clade, we calculated the fraction of PNP-encoding genes found in the genomic neighborhood of known anti-phage systems (where the genomic neighborhood was defined as 10 genes on each side of the PNP-encoding gene). On the other hand, the fraction of defense-associated genes expected by chance was computed for each clade by dividing the total number of genes located in the neighborhood of anti-phage systems by the total number of genes in the genomes of origin. These values were used to assess whether proteins from each clade colocalize with anti-phage systems more frequently than expected by chance, using a binomial test and FDR correction of the p-value (**Table S3**).

PNP domain sequences from the Cap17 clade were collected and a new tree was computed as described above, using the PNP domain of *E. coli* MtnN (IMG gene ID 2519179855) as an outgroup (**Fig. 2B**). Members of this clade were manually inspected for known genetic architectures of anti-phage systems.

To investigate the presence of ATP nucleosidases in eukaryotes, 10,176 eukaryotic proteins harboring a PNP domain (pfam PF01048) were collected from the Pfam database web page^73^. These proteins were grouped in 2,571 clusters based on sequence homology and searched for protein domains as described above. Similarly, cluster representatives with a hit to pfam PF01048 with an e-value equal to or lower than 0.01 and spanning at least 100 residues were selected and the sequence of the PNP domain was extracted. Prokaryotic and eukaryotic PNP sequences were aligned and a tree was computed as described above. Node support was computed using 5000 iterations of the ultrafast bootstrap function in IQtree (option -bb 5000)^42^. Arthropod and nematode proteins from the Cap17 clade with a Zα domain were expanded to the model species *D. melanogaster* (CG12065) and *C. elegans* (H14E04.2) by blastp search against the Uniprot database.

### Detection of Detocs in prokaryotic genomes

Homologs of DtcA, DtcB and DtcC were collected from the above-described protein clusters. For each protein, homologs were aligned using Clustal-omega^75^ and the resulting multi-sequence alignment was used to build an HMM profile using the hmmbuild function of hmmer (version 3.3.2)^78^. MacSyFinder^79^ was then used to search Detocs using the three HMM profiles, requiring all three genes to be present with up to one foreign gene between each Detocs gene. The resulting detected systems were manually inspected for false positives and classified by effector type.

## Supplementary Materials

**Table S1 – Anti-phage systems studied here**

**Table S2 – Phages used in this study**

**Table S3 – Genomic association of prokaryotic PNP homologs with known anti-phage systems**

**Table S4 – Prokaryotic PNP homologs from the Cap17 clade**

**Table S5 – Detection of Detocs in bacterial genomes**

**Table S6 – Eukaryotic homologs from the defensive PNP clade**

**Table S7 – Primers used for cloning point mutations**

**Table S8 – Sequences of purified proteins**

**Figure S1.**
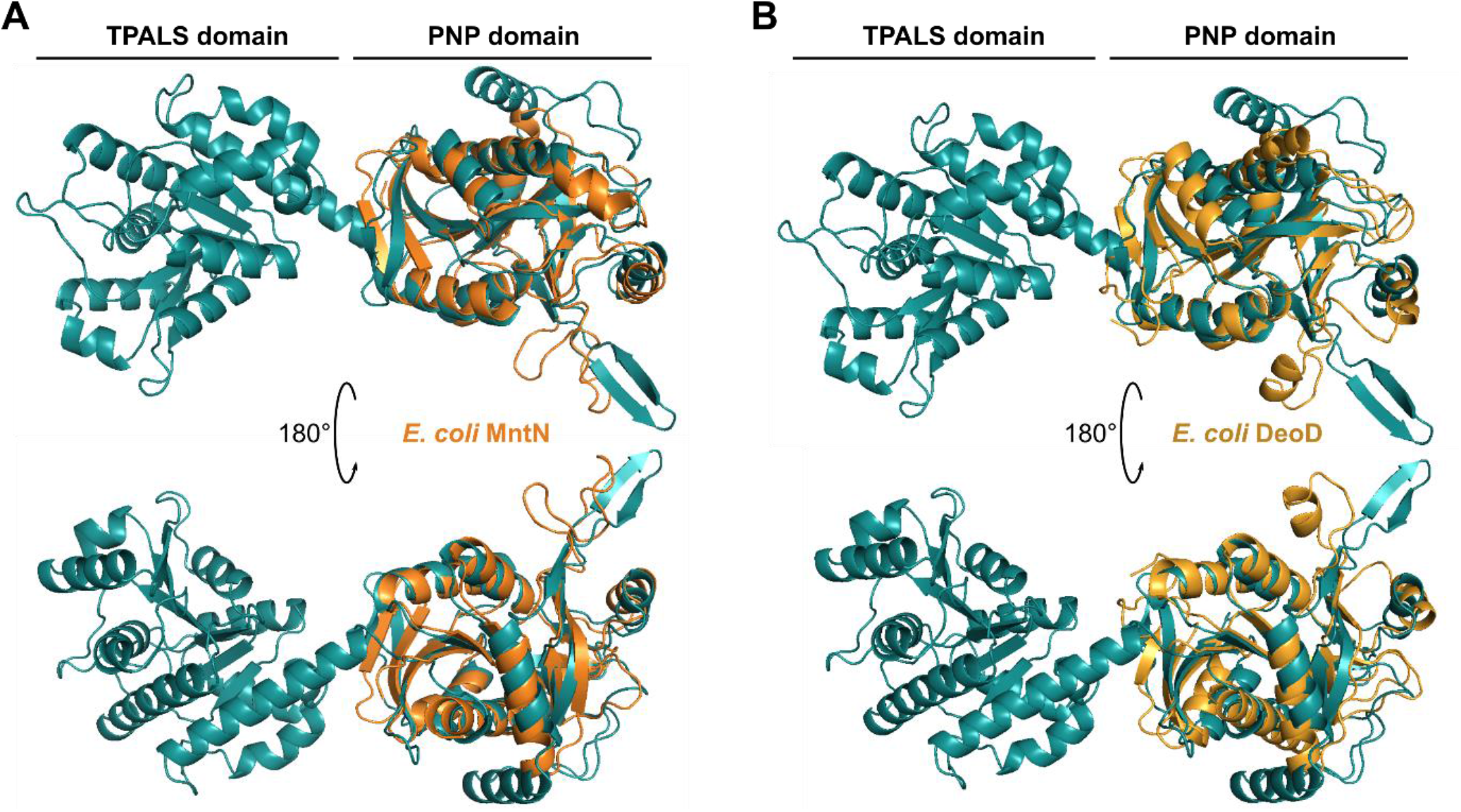
Structural homology of the Cap17 PNP domain with housekeeping PNP enzymes. The structure of *E. coli* Cap17 was predicted using AlphaFold^27,28^ and aligned to the experimentally-determined structures of (A) MtnN (PDB: 1NC1) and (B) DeoD (PDB: 1ECP) using the PDBeFold server^80^. Abbreviations: TPALS, TIR- and PNP-associating SLOG family; PNP, purine nucleoside phosphorylase.

**Figure S2.**
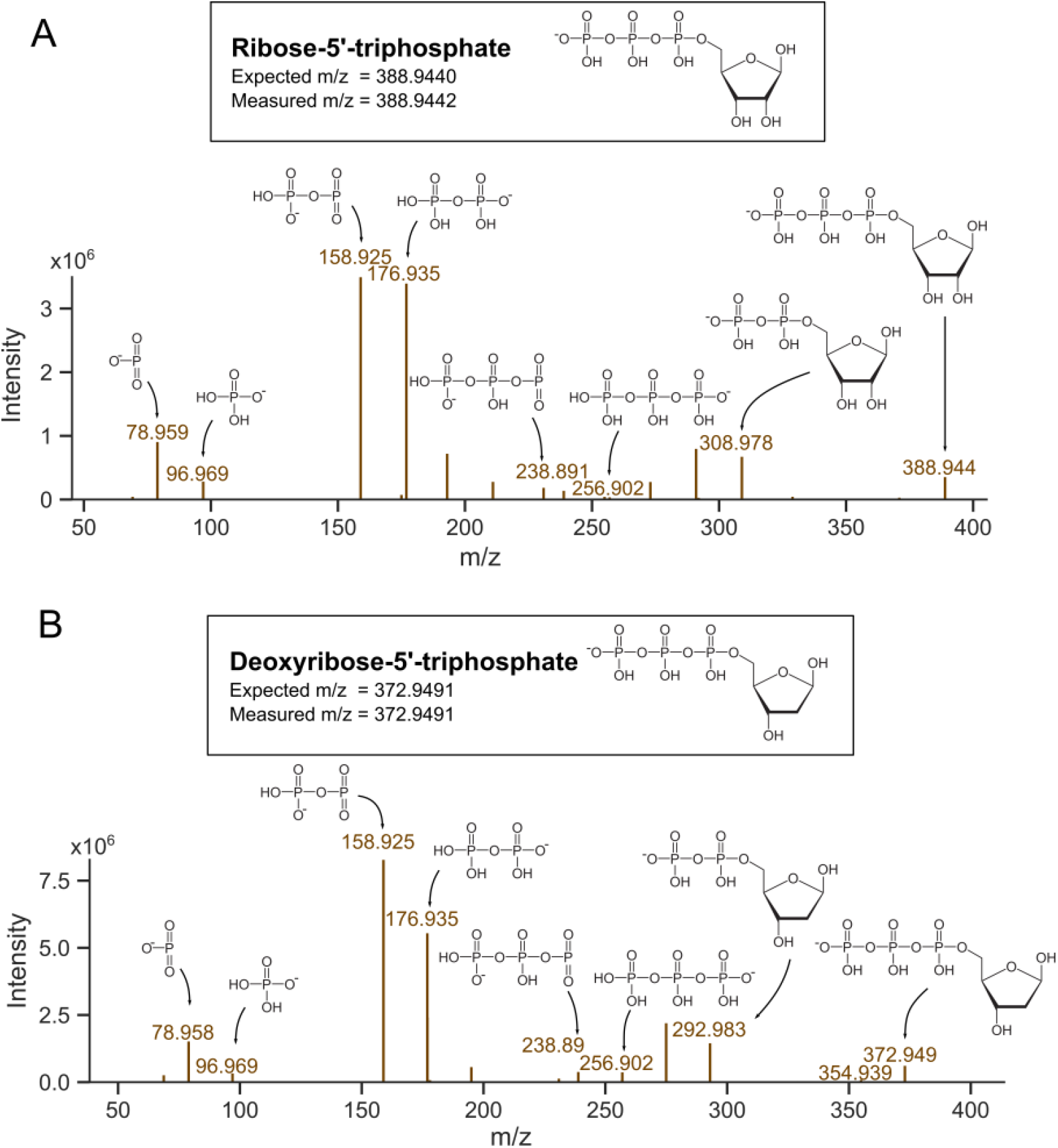
MS/MS fragmentation spectra. (A) The peak of m/z = 388.9442, a product of ATP conversion by Cap17 (**Fig. 1D**), was subjected to MS/MS, supporting the identification of this product as ribose-5’-triphosphate. (B) The peak of m/z = 372.9491, a product of dATP conversion by Cap17 (**Fig. 1E**) was subjected to MS/MS, supporting the identification of this product as deoxyribose-5’-triphosphate.

**Figure S3.**
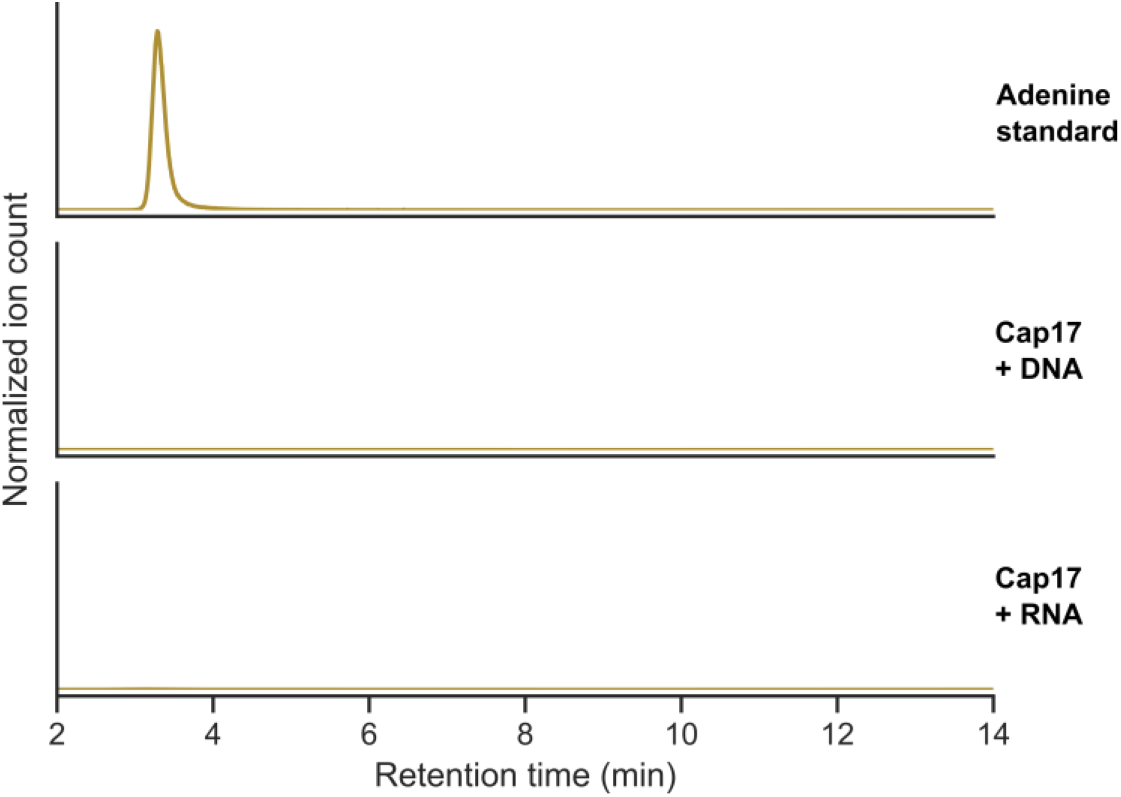
Cap17 is inactive on DNA and RNA. Purified Cap17 was incubated with DNA or RNA and reactions were analyzed by LC-MS. Mass chromatograms of ions with the mass of adenine (m/z = 134.0473) are shown, together with an adenine standard. Representative of three replicates.

**Fig. S4.**
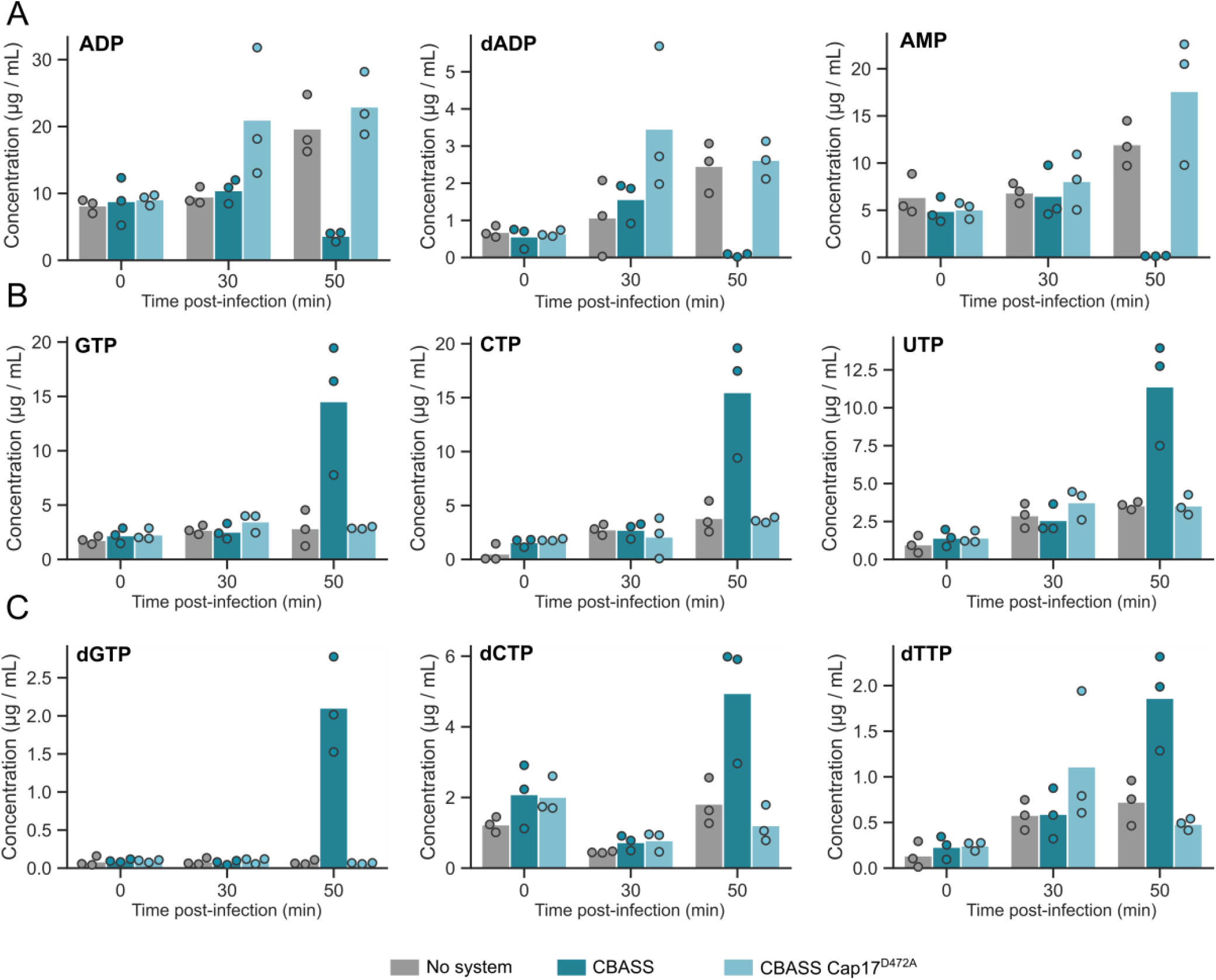
Nucleotide concentrations during P1 infection. Quantification of nucleotides in lysates derived from P1-infected *E. coli* K-12 cells expressing an empty vector (no system), a wild-type CBASS, or mutated CBASS (CBASS-Cap17^D472A^) by LC-MS. Bars represent the average of three replicates with individual data points overlaid.

**Figure S5.**
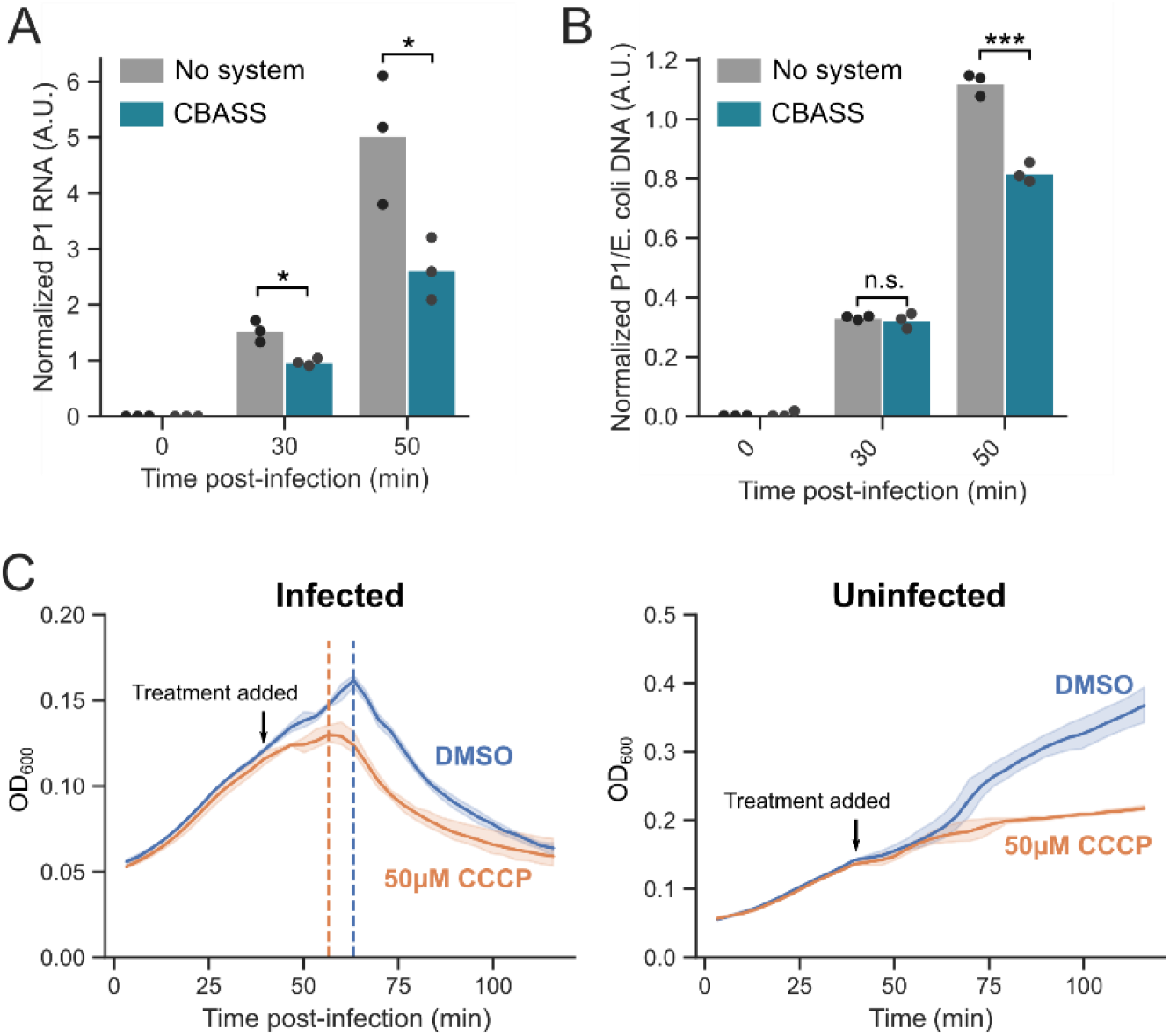
Pleiotropic effects of Cap17 activity during P1 infection. (A) RNA from P1-infected *E. coli* cells expressing a control vector or the CBASS system was sequenced. The number of phage reads normalized to reads from spiked-in *B. subtilis* RNA is shown. (B) DNA from P1-infected *E. coli* cells expressing a control vector or the CBASS system was sequenced. The ratio of phage to *E. coli* reads after normalization to reads from spiked-in *B. subtilis* DNA is shown. For panel (A) and (B), bars show the mean of three independent replicates with individual data points overlaid. Stars show significance levels of a two-sided t-test (*, p < 0.05; ***, p < 0.001; n.s., not significant). (C) *E. coli* cells were infected with phage P1 at MOI of 3 (left) or were left uninfected (right), and DMSO or 50 μM cyanide 3-chlorophenylhydrazone (CCCP) were added after 40 min. Growth was followed by optical density at 600 nm. Curves show the mean of three replicates with the standard deviation shown as a shaded area. Dashed lines represent temporal onset of lysis.

**Figure S6.**
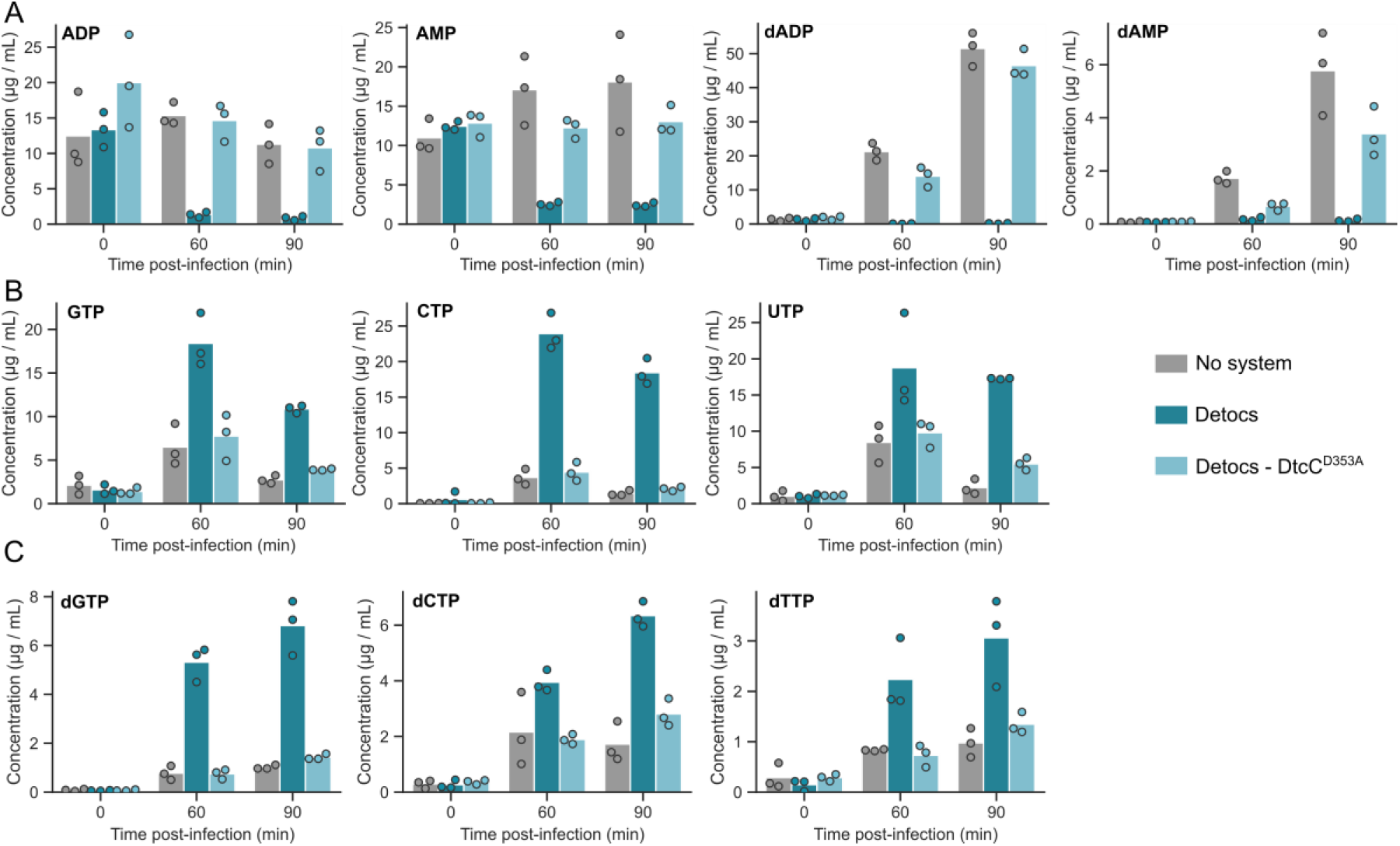
Nucleotide concentrations during T5 infection. Quantification of nucleotides in lysates derived from T5-infected *E. coli* cells expressing an empty vector (no system), a wild-type Detocs, or mutated Detocs (Detocs-DtcC^D353A^). Mean of three replicates, with individual data points overlaid.

**Figure S7.**
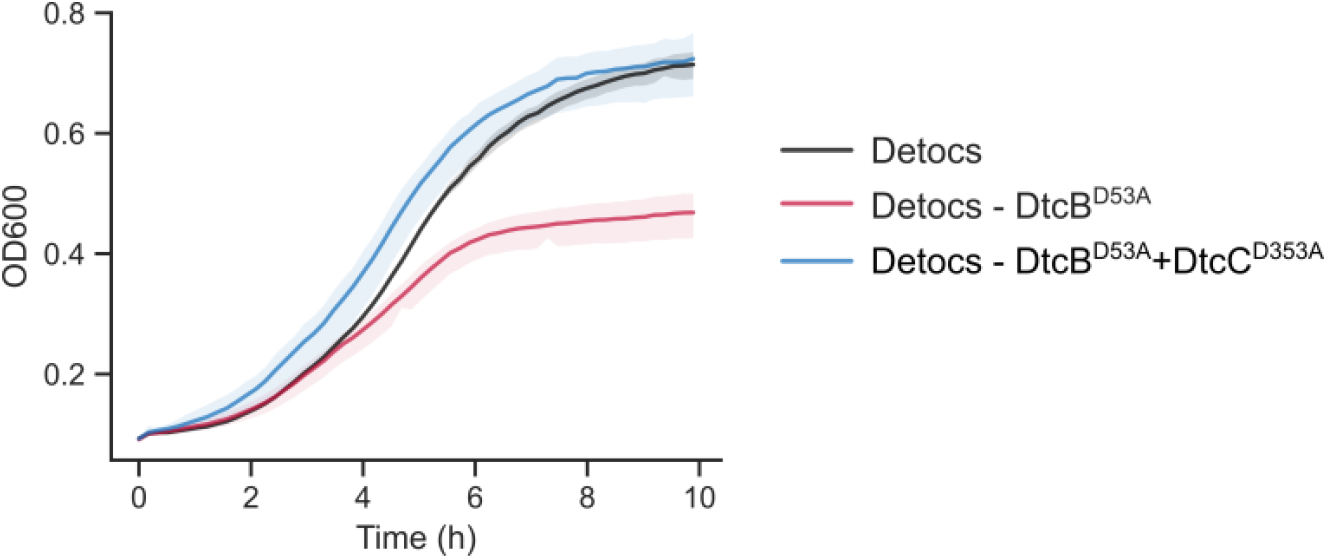
A point mutation in the receiver domain of DtcB is toxic in a PNP-dependent manner. Growth curves of *E. coli* cells expressing a wild-type Detocs from *V. alginolyticus* UCD-32C, a Detocs with a point mutation in the phosphate-receiving aspartate of DtcB, or a Detocs with this mutation combined with a point mutation in the PNP catalytic site of DtcC, as measured by optical density at 600 nm. Curves show the mean of three replicates with the standard deviation shown as a shaded area.

**Figure S8.**
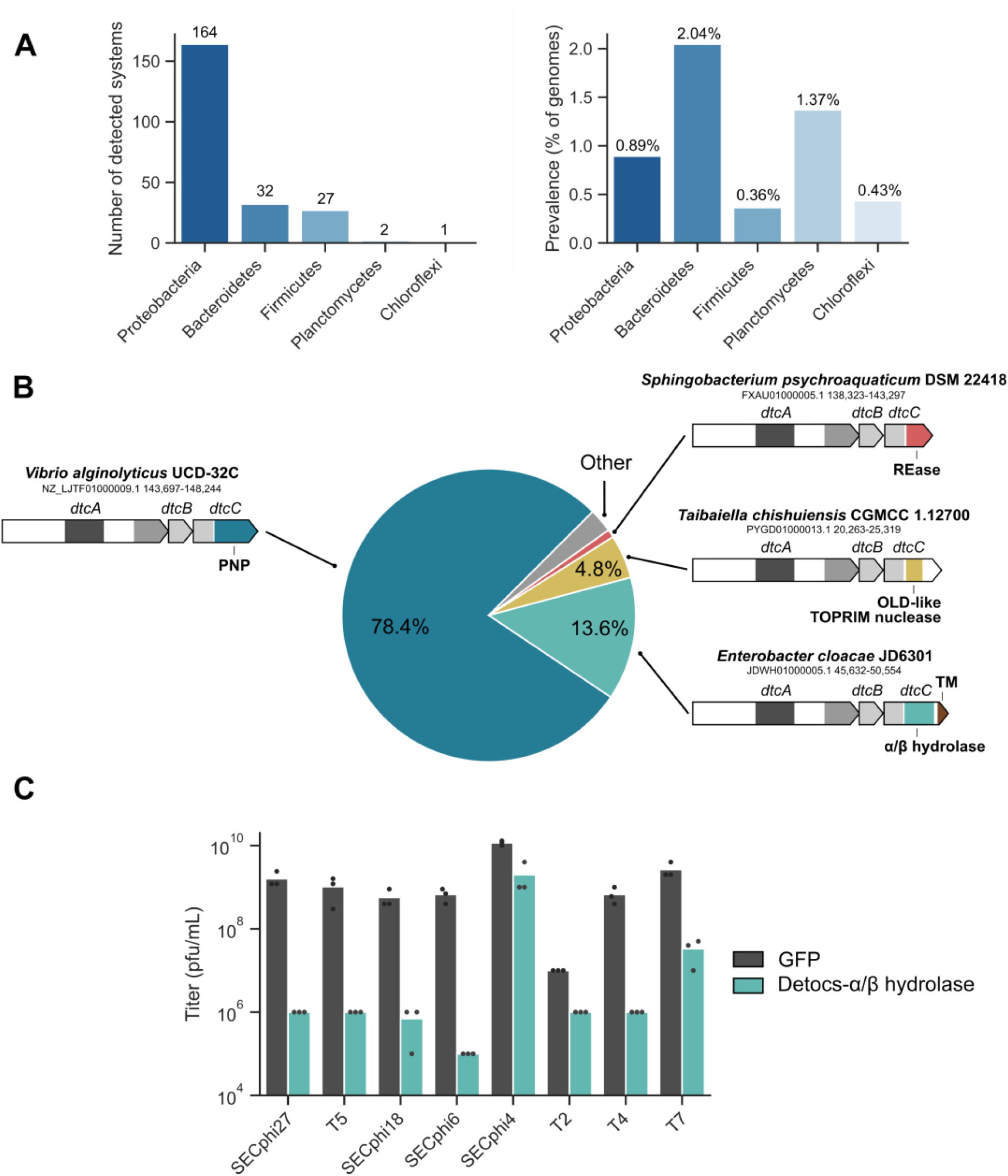
Detocs in the bacterial pangenome. (A) Detection of Detocs in bacterial genomes. For phyla in which Detocs is found, the number of systems (left) and the percentage of genomes carrying Detocs (right) are shown. (B) Distribution of types of effector domains in Detocs. Examples of systems are shown, with accession numbers and genome coordinates provided. Abbreviations: REase, Restriction Endonuclease (pfam PF18742); TOPRIM, Topoisomerase-Primase (pfam PF20469); TM, Transmembrane. (C) A Detocs system from *Enterobacter cloacae* with a transmembrane α/β hydrolase effector provides broad phage resistance in *E. coli*.

